# Endurance exercise ameliorates phenotypes in *Drosophila* models of Spinocerebellar Ataxias

**DOI:** 10.1101/2021.12.06.471345

**Authors:** Alyson Sujkowski, Kristin Richardson, Matthew V. Prifti, R. J. Wessells, Sokol V. Todi

**Affiliations:** Department of Physiology, Wayne State University School of Medicine; Department of Pharmacology, Wayne State University School of Medicine; Department of Neurology, Wayne State University School of Medicine

**Author notes:** Correspondence: 540 E Canfield, Scott Hall Rm 3108, Detroit, MI 48201, USA. Correspondence: 540 E Canfield, Scott Hall Rm 5275, Detroit, MI 48201, USA.

## Abstract

Endurance exercise is a potent intervention with widespread benefits proven to reduce disease incidence and impact across species. While endurance exercise supports neural plasticity, enhanced memory, and reduced neurodegeneration, less is known about the effect of chronic exercise on the progression of movement disorders such as ataxias. Here, we focused on three different types of ataxias, Spinocerebellar Ataxias Type (SCAs) 2, 3, and 6, belonging to the polyglutamine (polyQ) family of neurodegenerative disorders. In *Drosophila* models of these SCAs, flies progressively lose motor function. Here, we observe marked protection of speed and endurance in exercised SCA2 flies and modest protection in exercised SCA6 models, while no benefit is observed in SCA3 flies. Causative protein levels are reduced in SCA2 flies after chronic exercise, but not in SCA3 models, linking protein levels to exercise-based benefits. Additional investigations indicate that the exercise-inducible protein, Sestrin (Sesn) suppresses mobility decline and improves early death in SCA2 flies, even without exercise, coincident with disease protein level reduction and increased autophagic flux. These improvements depend on previously established functions of Sesn that reduce oxidative damage and modulate mTOR activity. Our study suggests differential responses of polyQ SCAs to exercise, highlighting the potential for more extensive application of exercise-based therapies in the prevention of polyQ neurodegeneration. Defining the mechanisms by which endurance exercise suppresses polyQ SCAs will open the door for more effective treatment for these diseases.

## Introduction

Nine inherited neurodegenerative disorders are caused by expansion of a CAG triplet repeat in the protein-coding region of the respective disease genes. The CAG repeat encodes the amino acid glutamine; thus, the disease protein carries a lengthened tract of polyglutamines (polyQ) that renders it toxic. The polyQ family includes six Spinocerebellar Ataxias (SCAs 1, 2, 3, 6, 7, and 17), Huntington’s Disease, Dentatorubral-pallidoluysian Atrophy, and Kennedy’s disease ^(1-7)^. Various pathogenic mechanisms are shared, involving primarily toxic gain-of-function acquired by the disease protein. These mutations yield aggregation of the toxic polyQ protein and despite their wide expression throughout the body, only select central nervous system regions degenerate in each disease. Other common mechanisms underlying polyQ pathogenesis include mitochondrial, transcriptional, and autophagy dysfunction (8). Although the disorders share disease mechanisms, the individual host proteins have divergent functions (1–7).

There is an unmet need for the treatment of polyQ diseases as there is currently no therapeutic solution for any of these disorders. Here, we examined the idea that daily endurance exercise might be of benefit against polyQ disorders. Endurance exercise is a low-cost, effective intervention with broad pro-healthspan effects, including reducing the incidence of obesity, heart disease, cancer, and cognitive defects (9–11). Previous studies examined to a limited extent the effects of chronic exercise in patients with advanced stages of ataxia, including polyQ disorders: some types of exercise might help in certain cases (12–14) but not others (15, 16). These studies may have been limited by lack of normalized protocols, genetics, and variable disease stage among patients. Some work on the possible role of exercise was also conducted in mouse models of HD, SBMA and SCA1. SCA1 and SBMA mice seemed to benefit from exercise (17–20), whereas some models of HD improved (21, 22) while others did not (23).

Differences in these studies could be due to variation in protocols, genetic background, age of testing and other parameters difficult to normalize among studies. While studies in animal models have identified potential pathways by which exercise may affect polyQ phenotypes, absence of normalization may limit the overall applicability and extension of those results to other polyQ diseases and further up the evolutionary chain. Here, we take advantage of the *Drosophila* model system to systematically examine which polyQ disorders are responsive to exercise therapy and to identify mechanisms involved.

*Drosophila melanogaster* has been widely and successfully utilized to understand polyQ disease mechanisms (24–27). In parallel, the fly has emerged as a highly efficient model for studying long-term effects of chronic exercise (28). Following a 3-week daily, gradually ramped training program, flies of diverse genotypes experience reproducible changes to speed, endurance, flight, cardiac performance, mitophagy, and mitochondrial function (29–36). These changes track with adaptive physiological changes seen in chronically exercising humans (28, 37), and induce conserved changes in gene expression, including conserved mediators of exercise benefits, such as PGC1-a and Sesn (38, 39).

Here, we tested the effects of chronic exercise in three *Drosophila* models of polyQ SCAs: 2, 3, and 6. Ectopic expression of these human, polyQ-expanded disease proteins in flies is toxic (40–47). We find that exercise has a dramatically positive effect in SCA2 flies, with modest effects on SCA6 model flies and no benefit to SCA3 flies, indicating a range of response in different disease models. In addition, the exercise-mimicking protein, Sesn improves early death and rescues mobility declines in SCA2 model flies. We further find that improvement in disease phenotypes depends on functions of Sesn that reduce oxidative damage and modulate mTOR. Our observations identify Sesn as a key genetic mediator underlying the protective effects of exercise and highlight mechanistic targets that can be leveraged towards therapeutic options for patients unable to exercise, or to supplement the benefits of those who can.

## Results

### Exercise differentially impacts mobility in *Drosophila* polyQ SCA models

Expression of polyQ disease proteins in flies causes reduced motility and neuronal cell death, as we and others have shown in various publications (24-27, 41, 45-52). To test if endurance exercise reduces polyQ-dependent phenotypes in the fly, we selected three SCAs: 2, 3, and 6 (Figure 1A). Full-length, human disease protein was expressed through the binary, RU486-inducible Gal4-UAS system (32, 53) in all fly neurons and only during adulthood, which is toxic (40–47). We have established an optimized exercise protocol (Figure 1B) that drives reproducible effects that are robust across genotypes (29, 34). These effects are not due to non-specific stress or behavioral effect of the regimen—unexercised flies exposed to training, but prevented from running, do not experience improvements (29, 34).

**Figure 1:**
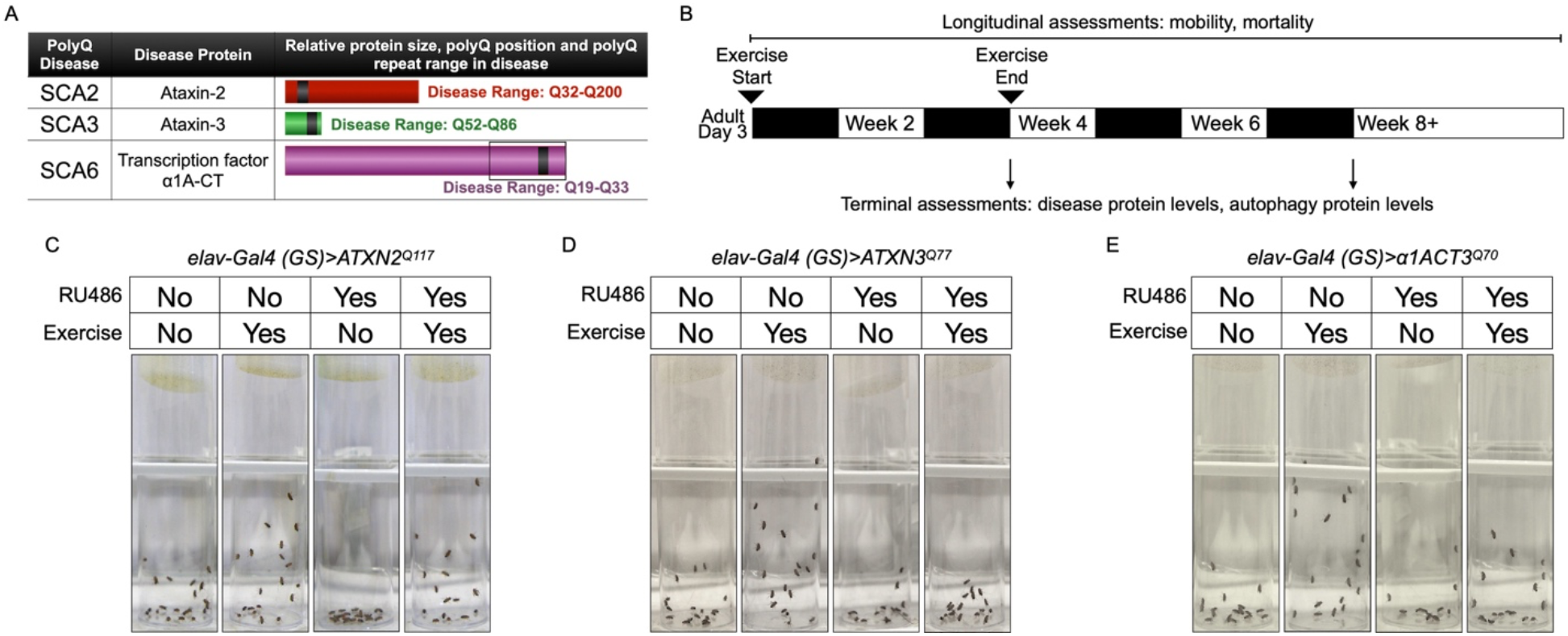
Endurance exercise differentially affects mobility in *Drosophila* models of Spinocerebellar Ataxia. **(A)** Table of Spinocerebellar Ataxia (SCA) models used in this study. **(B)** Timeline of endurance exercise program and assessment of physiology and disease protein levels. **(C-E)** Representative climbing speed images in *Drosophila* models of **(C)** SCA2, **(D)** SCA3 and **(E)** SCA6. Photos taken 2 seconds after inducing negative geotaxis response in 4-week-old flies, after endurance exercise is complete.

Exercised wild-type *Drosophila* retain faster climbing speed than age-matched, unexercised siblings (28). SCA2 model flies expressing polyQ-expanded ATXN2 specifically in adult neurons (*elav-Gal4 (GS)>ATXN2^Q117^*) have reduced climbing speed by 4 weeks of age, but exercise fully rescues climbing speed to the level of age-matched, uninduced control flies that complete our three-week, ramped exercise program (Figure 1C, Supplementary video S1). In contrast, four-week-old SCA3 model flies (*elav-Gal4 (GS)>ATXN3^Q77^*) have lower climbing speed than unexercised, uninduced control flies and do not increase climbing speed after exercise training (Figure 1D). Flies expressing hyper-expanded α1ACT (*elav-Gal4 (GS)>α1ACTQ^70^*), the disease gene in SCA6 (44), also have reduced week-four climbing speed in comparison to uninduced, unexercised control flies, but climbing speed is partially rescued after exercise training (Figure 1E).

PolyQ disorders are progressive and neurodegenerative severity tends to increase with age (1, 2). We have previously shown that exercise-induced improvements to climbing speed and endurance are long-lasting and persist even after the three-week exercise program is complete (29). Neither RU486 feeding nor expression of the empty vector used to generate transgenic flies in adult neurons negatively affect exercise adaptations to either climbing speed or endurance. Furthermore, expression of an essentially pure polyQ80 repeat (54) in adult neurons reduced climbing speed and abrogated exercise benefits, while age-matched, uninduced control flies adapted to exercise normally (Supplementary Figure S1A-C).

Having confirmed that post-exercise differences in climbing speed can be discerned in polyQ model flies, we next subjected the same three SCA models from Figure 1 to our endurance exercise program and tested them longitudinally for changes to climbing speed and endurance. In contrast to the ramped exercise program, endurance is tested by placing flies on the climbing apparatus on non-training days and allowing them to climb to exhaustion. When fewer than 10% of flies in a vial no longer respond to the climbing stimulus, the time is recorded, and the vial is removed from the apparatus and scored as fatigued. These data are plotted similarly to a survival curve, with each datum representing a single vial of 20 flies (55).

SCA2 model flies have reduced climbing speed by adult week four (Figure 2A, compare RU-UN to RU+UN), but endurance exercise fully rescues climbing speed to the level of exercised, uninduced control flies, and improvement persists into adult week five (Figure 2A, compare RU-EX to RU+EX). Endurance in SCA2 model flies is similar to uninduced, unexercised siblings (log-rank, p=0.2018) and exercised SCA2 model flies improve endurance as well as exercised, uninduced controls (Figure 2B). We next examined ATXN2 protein levels to see if improved climbing correlated with changes in toxic protein levels. Exercise markedly reduced ATXN2 levels relative to age-matched, unexercised siblings, linking improved physiology to reductions in disease protein (Figure 1C).

**Figure 2:**
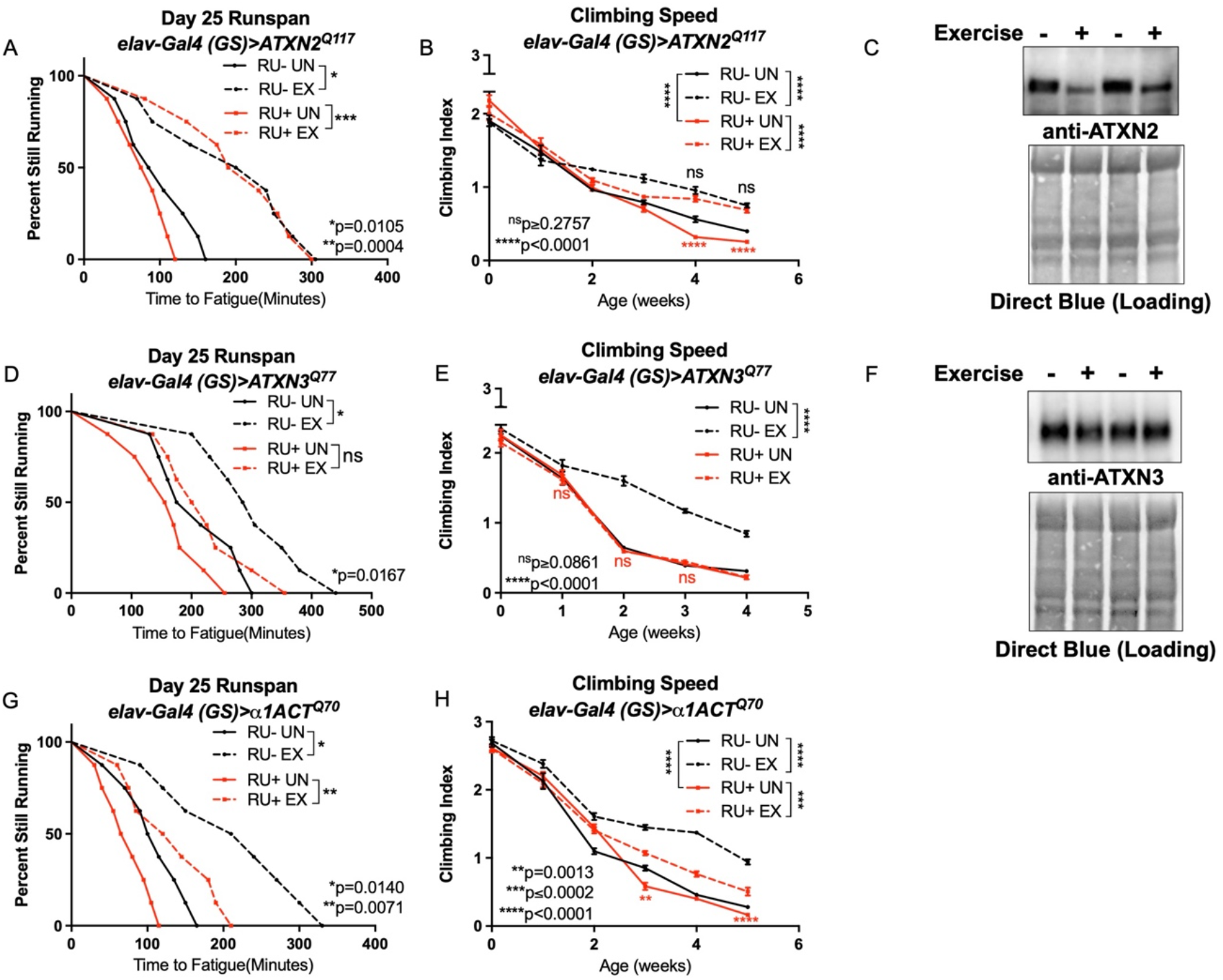
Endurance exercise differentially affects mobility and disease protein levels *Drosophila* models of SCA. **(A)** Flies ectopically expressing polyQ-expanded *ATXN2* in adult neurons (RU+) have lower climbing speed than unexercised, uninduced control flies (RU-UN) by adult week 4. Exercise fully rescues climbing speed to the level of exercised, uninduced control flies (compare RU-EX to RU+ EX). **(B)** Flies expressing polyQ-expanded *ATXN2* in adult neurons (RU+) have similar endurance to uninduced control flies (RU-) whether exercised or not. **(C)** Exercise reduces ATXN2 levels in flies ectopically expressing polyQ-expanded *ATXN2* in adult neurons. **(D-E)** Flies ectopically expressing polyQ-expanded *ATXN3* in adult neurons (RU+) have similar **(D)** climbing speed and **(E)** endurance to uninduced, unexercised control flies and fail to improve either **(D)** climbing speed or **(E)** endurance with exercise. **(F)** Exercise does not affect ATXN3 levels in flies expressing polyQ-expanded *ATXN3* in adult neurons. **(G)** Flies ectopically expressing polyQ-expanded *α1ACT* in adult neurons (RU+) have lower climbing speed than unexercised, uninduced control flies (RU-UN) by adult week 3, and exercise partially rescues climbing speed, although not to the level of exercised, uninduced control flies. **(H)** Exercise improves endurance in flies expressing polyQ-expanded *α1ACT* in adult neurons (RU+ EX), but endurance is not as good as exercised, uninduced control siblings (compare RU-EX to RU+ EX). Mobility and endurance experiments performed in triplicate. n≥100 (climbing speed), n≥8 vials of 20 flies (endurance). Error bars indicate +/-SD.

In contrast, SCA3 model flies do not improve climbing speed (Figure 2D) or endurance (Figure 2E) after exercise training. Furthermore, exercised and unexercised SCA3 flies had similar levels of ATXN3, the causative, polyQ-expanded disease protein in SCA3 (15) (Figure 2F). SCA6 model flies had reductions in climbing speed at weeks three and five compared to unexercised, uninduced siblings, and exercise training improved climbing speed, albeit not to the level of uninduced, exercised control flies (Figure 2G). Unexercised SCA6 flies trend toward lower endurance relative to uninduced, unexercised control flies (log-rank, p=0.0559) and improve endurance after exercise training, but do not equal the endurance of uninduced, exercised controls (log-rank, p=0.0218) (Figure 2H).

### Sestrin expression improves phenotypes in SCA2 model flies without exercise

We have previously shown that the stress-inducible protein Sesn is necessary for exercise adaptations in multiple species (39), and Sesn activity has been proposed to play a protective role against neurodegeneration (56). Both endurance exercise and Sesn increase autophagy by inhibiting mTOR (32, 57, 58), and increased autophagy increases ATXN2 solubility and reduces phenotypes in SCA2 patient cells (59). Since muscle-specific dSesn overexpression is sufficient to replicate the beneficial adaptations of endurance exercise, even in sedentary *Drosophila* (39), we next overexpressed dSesn in SCA2 model flies.

Longevity assessment in SCA model flies expressing polyQ-expanded disease protein selectively in adult neurons confirmed previously reported lifespan reductions in both SCA3 and SCA6 model flies (44, 60) (Figure 3B, C), but neither ATXN2 expression nor exercise significantly affected longevity in SCA2 flies (Figure 3A). *ATXN2* CAG expansion causes early mortality in SCA2 patients (61); therefore, we performed subsequent experiments in flies ubiquitously expressing two copies of polyQ-expanded ATXN2 simultaneously with dSesn overexpression (*sqh>ATXN2^Q117^;dSesn^WT^;ATXN2^Q117^*) to more closely model clinical subjects. Indeed, constitutive expression of polyQ-expanded ATXN2 in all tissues, throughout development and in adults, (*sqh>ATXN2^Q117^;ATXN2^Q117^*) caused early death in both female (Figure 4A) and male (Figure 4D) flies. Mobility reductions appeared earlier than with pan-neuronal expression, present upon adult eclosion from the pupal case and persisting until death (Figure 4B, E). On the other hand, flies ubiquitously expressing both ATXN2^Q117^ and dSesn improved survival (Figure 3A, D), although rescue was not to the level of background control flies. Mobility improvements were more pronounced, with complete rescue of climbing speed in both females (Figure 3B) and males (Figure 3E). We next examined disease protein levels in *sqh>ATXN2^Q117^;dSesn^WT^;ATXN2^Q117^* flies and found significant reduction in ATXN2 compared to SCA2 flies without dSesn expression (Figure 4C, F, quantified in Figure 7), similar to aforementioned reduction in disease protein after exercise. In contrast, ubiquitous expression of either ATXN3^Q77^ or α1ACT^Q70^ leads to developmental and early adult lethality (44, 60), and dSesn expression did not improve outcomes (data not shown).

**Figure 3:**
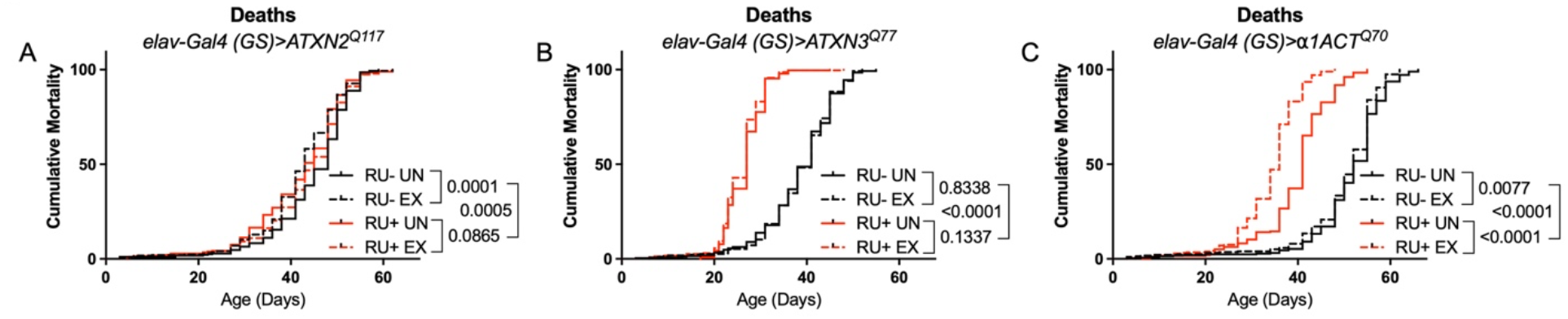
Expression of Atxn2 in adult neurons does not negatively impact lifespan. Cumulative mortality in **(A)** *elav-Gal4 (GS)>ATXN2^Q117^* flies is similar whether exercised or not. **(B)** *elav-Gal4 (GS)>ATXN3^Q77^* flies have reduced lifespan compared to uninduced controls, and exercise does not negatively affect either group. **(C)** *elav-Gal4 (GS)>α1ACT^Q70^* flies have lower lifespan than uninduced controls, and exercise reduces lifespan further. n≥231.

**Figure 4:**
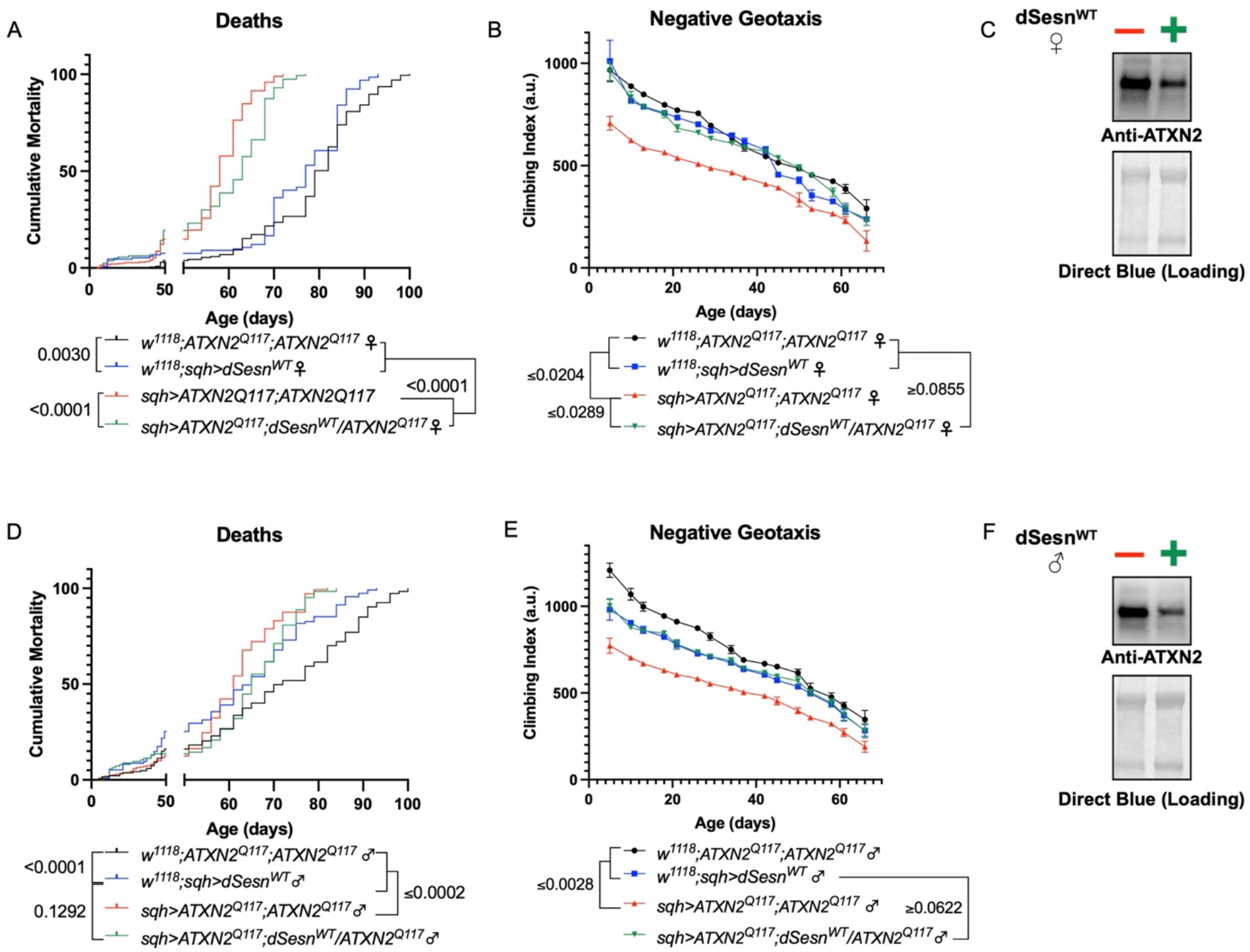
dSesn expression improves early death and low mobility in SCA2 flies, concurrent with reductions in disease protein. Female **(A, B)** and male **(D, E)** flies ubiquitously expressing polyQ-expanded ATXN2 (red lines) have early death **(A, D)** and lower climbing speed **(B, E)** than age-matched background control flies (blue and black lines). dSesn expression in flies also expressing polyQ-expanded ATXN2 (green lines) partially rescues early death **(A, D)** and fully rescues decreased mobility **(B, E).** ATXN2 levels are lower in both female **(C)** and male **(F)** flies ubiquitously expressing both dSesn and polyQ-expanded ATXN2. Survival and mobility experiments performed in triplicate. n≥200 (survival). n≥100 (climbing speed). Error bars indicate +/-SD. Representative Western blots from four biological repetitions, quantified in Figure 6.

### Oxidative resistance and mTOR modulating functions of Sestrin are necessary to improve SCA2 phenotypes

Sestrin is a multi-functional protein with separate, previously identified oxidoreductase and mTOR interacting domains (62). The C86S (dSesn^CS^) mutation abolishes oxidoreductase activity while the D424A (dSesn^DA^) and D423A/D424A (dSesn^DDAA^) substitutions disrupt TORC1 inhibiting and TORC2/Akt potentiating functions of dSesn (62). We previously showed that interaction with mTOR is required for the mobility-extending effects of dSesn in wild-type, sedentary flies (39). Overexpression of either wild-type or mutated dSesns does not increase longevity in a wild-type background (i.e. without polyQ protein expression; Supplementary Figure S2). To explore the requirement of Sesn’s various functions for suppression of polyQ phenotypes, we performed mobility, survival and protein expression experiments in which we ubiquitously overexpressed dSesn^CS^, dSesn^DA^, dSesn^DDAA^, or wild-type dSesn (dSesn^WT^) in SCA2 flies.

First, we overexpressed dSesn without its antioxidant function (*sqh>ATXN2^Q117^;dSesn^CS^;ATXN2^Q117^*). Ubiquitous expression of dSesn^CS^ in SCA2 model flies does not provide any protection—in fact, it exacerbates premature death in both male and female flies (Figure 5A, D). However, we observed pronounced mobility improvements in female flies (Figure 5B) and mobility is fully rescued in males (Figure 5E). Whereas wild-type dSesn overexpression reduces ATXN2 levels in both female (Figure 5C) and male (Figure 5F) SCA2 model flies, dSesn^CS^ overexpression does not have the same impact (Figure 5C, F, quantified in Figure 7). These results suggest that while the oxidoreductase function of dSesn is not required for protection against motility defects in SCA2 flies, it is essential for improved survival.

**Figure 5:**
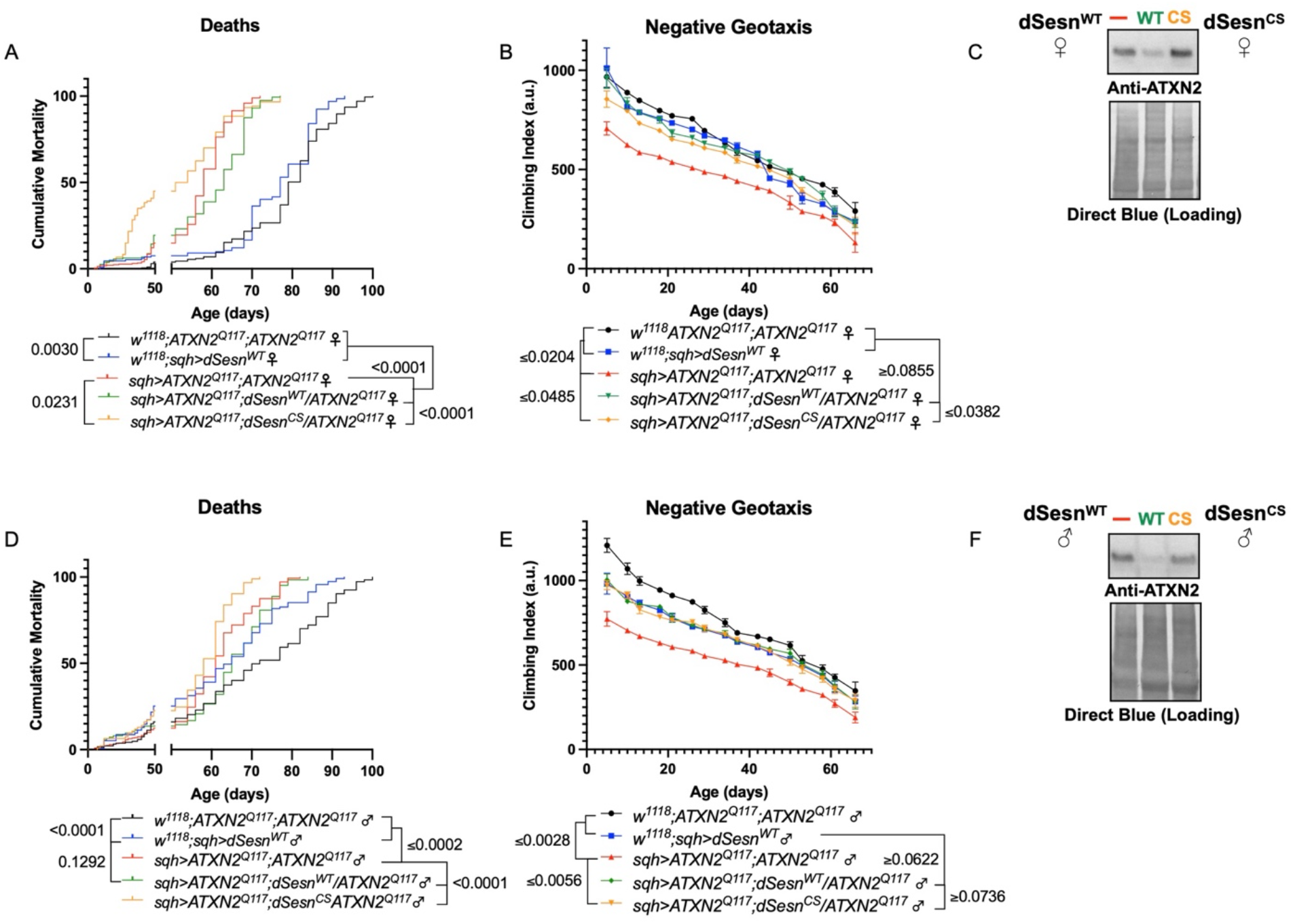
Oxidoreductase function is dispensable for mobility-extending effects of dSesn in flies expressing polyQ-expanded ATXN2. Ubiquitous expression of dSesn harboring a point mutation that abolishes oxidoreductase activity (*dSesn^CS^*, orange lines) exacerbates early death in SCA2 model flies (compare orange lines to red lines) in both females **(A)** and males **(D)**. In contrast, climbing speed is partially rescued in the same female flies (expressing both dSesn^CS^ and polyQ-expanded ATXN2, in orange) **(B)** and fully rescued in males **(E)**. **(C, F)** Wild-type dSesn expression reduces ATXN2 levels, but dSesn^CS^ expression does not. Survival and mobility experiments performed in triplicate. n≥200 (survival). n≥100 (climbing speed). Survival and climbing speed graphs depicted from same repetition depicted in Figure 2. Error bars indicate +/-SD. Representative Western blots from four biological repetitions, quantified in Figure 6.

We next ubiquitously expressed two separate amino-acid substitutions of dSesn that knock out its ability to modulate mTOR (62). Like dSesn^CS^ experiments, expression of either dSesn^DA^ (Figure 6A-D) or dSesn^DDAA^ (Figure 6E-H) increases early death in both female (Figure 6A, E) and male (Figure 6C, G) SCA2 model flies. In female flies, both dSesn^DA^ and dSesn^DDAA^ confer modest improvements to climbing speed compared to SCA2 model flies, but improvement does not equal that of wild type dSesn expression and is absent at later timepoints (Figure 6B, F). Male flies ubiquitously expressing dSesn^DA^ does not rescue mobility defects at any age (Figure 6D), while overexpression of dSesn^DDAA^ provides modest climbing speed improvements at early ages only (Figure 6H). Thus, interaction with mTOR appears critical for full rescue of climbing speed and protection against early death in SCA2 model flies.

**Figure 6:**
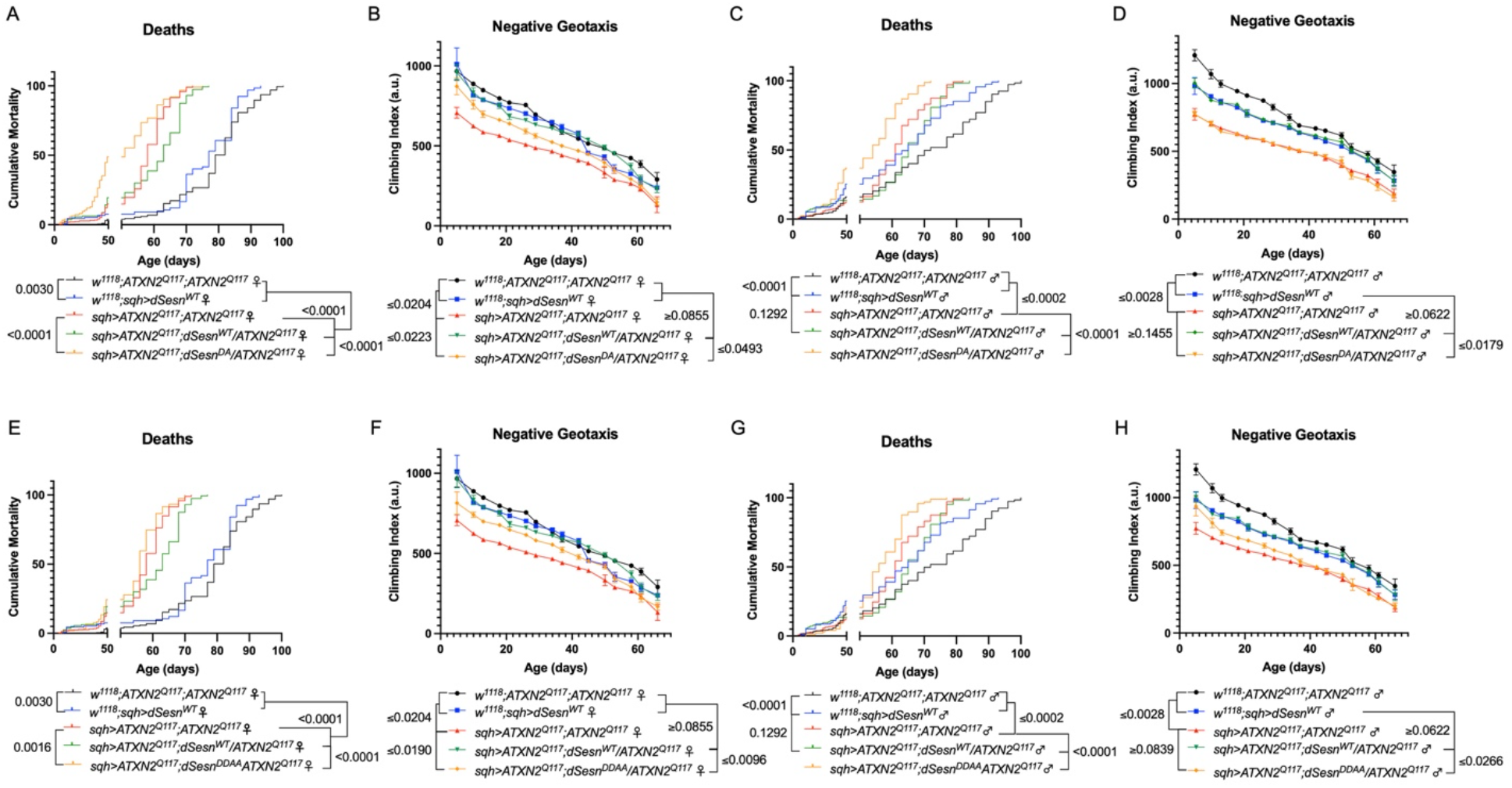
Interaction with mTOR is required for dSesn to improve survival and mobility in flies expressing polyQ-expanded ATXN2. Ubiquitous expression of dSesn harboring two separate mutations that abolish mTORC interaction *dSesn^DA^* **(A-D)**, *dSesn^DDAA^* **(E-H)** orange lines) exacerbates early death **(A,E,** females, **C,G,** males) and fails to rescue mobility **(B,F,** females, **D,H,** males) in SCA2 model flies. Survival and mobility experiments performed in triplicate. Survival and climbing speed graphs depicted from same repetition depicted in Figure 2. n≥200 (survival). n≥100 (climbing speed). Error bars indicate +/-SD.

To examine the relative impact of dSesn expression on disease protein levels, we again performed Western blots on flies ubiquitously expressing polyQ-expanded ATXN2 with and without simultaneous expression of either wild type dSesn or dSesn harboring mutations that separately abolish its oxidative protection or mTOR modulating functions. ATXN2 protein in SCA2 model flies (*sqh>ATXN2^Q117^;ATXN2^Q117^*) was easily visible by log-phase of mortality (Figure 7A-C, F). Although all dSesn-expressing cohorts had variable reductions in ATXN2 levels, only expression of wild-type dSesn correlated with significantly reduced disease protein. These data may reflect different mechanisms of protection at play and are discussed further below.

**Figure 7:**
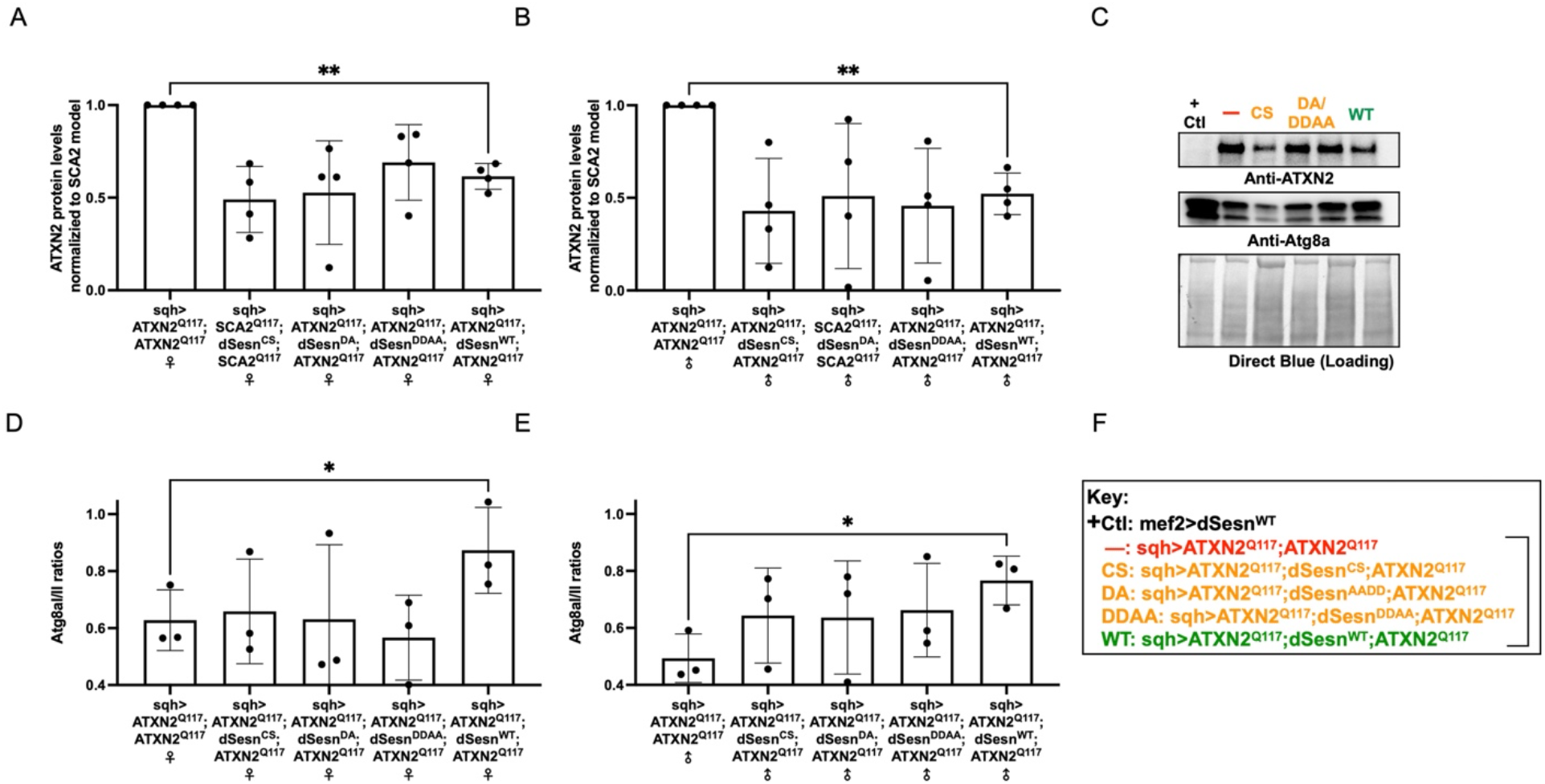
Wild-type dSesn expression reduces disease protein and increases autophagy in flies expressing polyQ-expanded ATXN2. Ubiquitous expression of dSesn significantly reduces ATXN2 levels **(A-C**, n=4) and increases Atg1a;Atg2a ratios **(C-E**, n=3). **(F)** Genotypes in representative Western blot and from independent biological replicates, quantified in A, B, D, and E. Select individual Western blots repetitions shown in previous Figures 4 and 5. Bracket indicates quantifications from A, B, D, and E.

Based on these results and on our prior observations in exercised flies and flies with muscle-specific dSesn overexpression (39, 63), we hypothesized that dSesn overexpression and reduced ATXN2 levels relate to changes in autophagy. We observed that AtgIa/IIa ratios increased in SCA2 model flies that were overexpressing wild-type dSesn, but not its mutated versions, which trend toward increased autophagy without reaching significance (Figure 7C, D-F).

Finally, since dSesn overexpression improved survival in flies ubiquitously expressing two copies of the polyQ-expanded ATXN2 transgene, we repeated our longevity experiments in this model of exercised SCA2 flies. Exercised female flies expressing two copies of polyQ-expanded ATXN2 either ubiquitously (Figure 8A) or in adult neurons (Figure 8C) significantly improved survival over unexercised siblings. Male SCA2 model flies trended toward improved survival in early life only, while they were still being subjected to exercise training (Figure 8B, D). Altogether, these findings indicate protective effects from endurance exercise not only in terms of motility (Figures 1, 3) but also in terms of longevity (Figure 8) in SCA2 flies.

**Figure 8:**
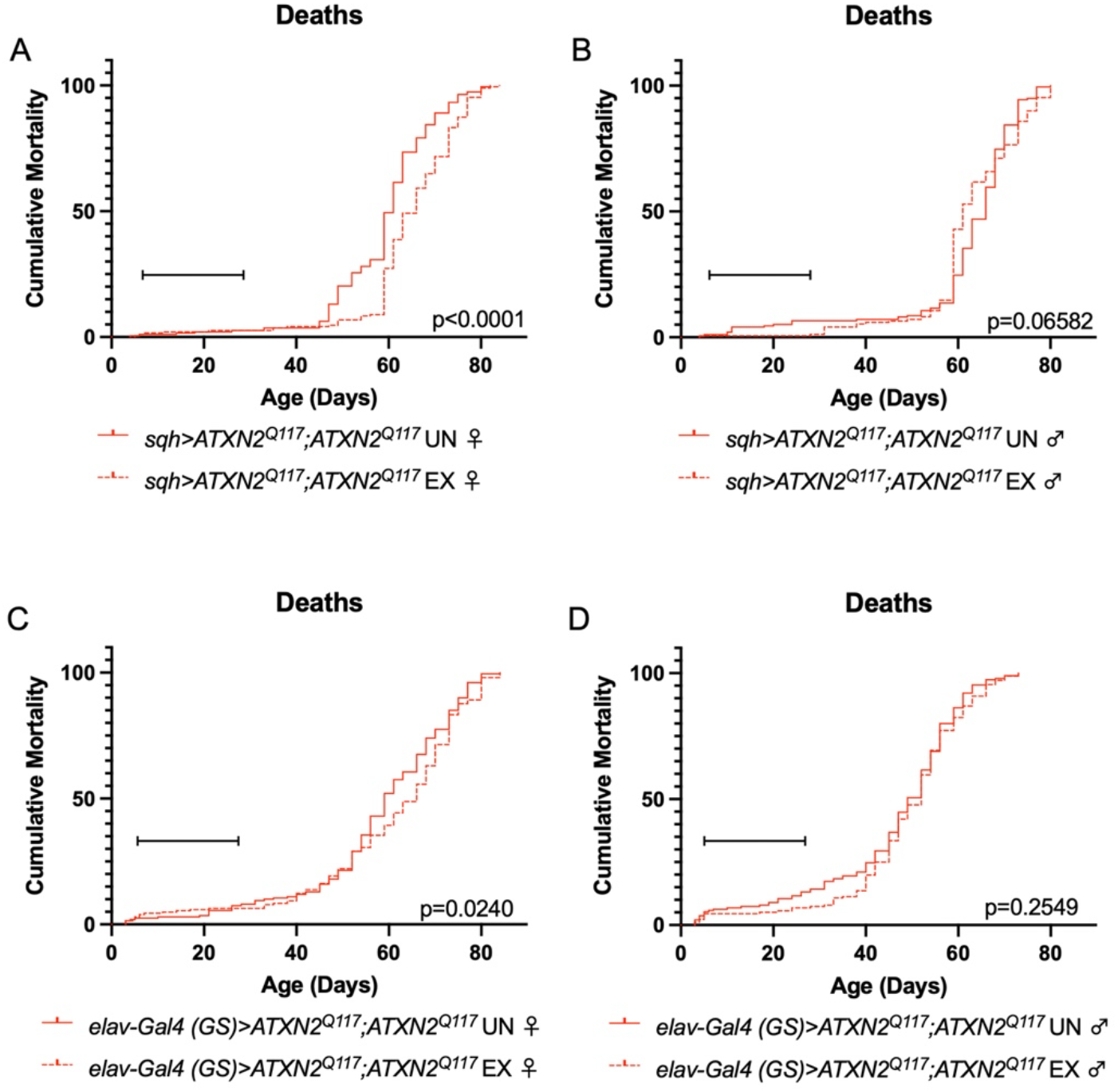
Exercise reduces early death in SCA2 model flies expressing two copies of ATXN2. Exercised female flies expressing two copies of polyQ expanded SCA2 ubiquitously **(A)** or in adult neurons **(C)** have increased survival compared to age-matched, unexercised siblings, while exercised male flies expressing two copies of polyQ expanded SCA2 ubiquitously **(B)** or in adult neurons **(D)** trend toward increased survival only in the first 25 days, the period in which flies are still undergoing exercise training. Brackets indicate exercise training period. P-values indicate log-rank for entire survival curve, n≥170, performed in duplicate.

**Figure 9:**
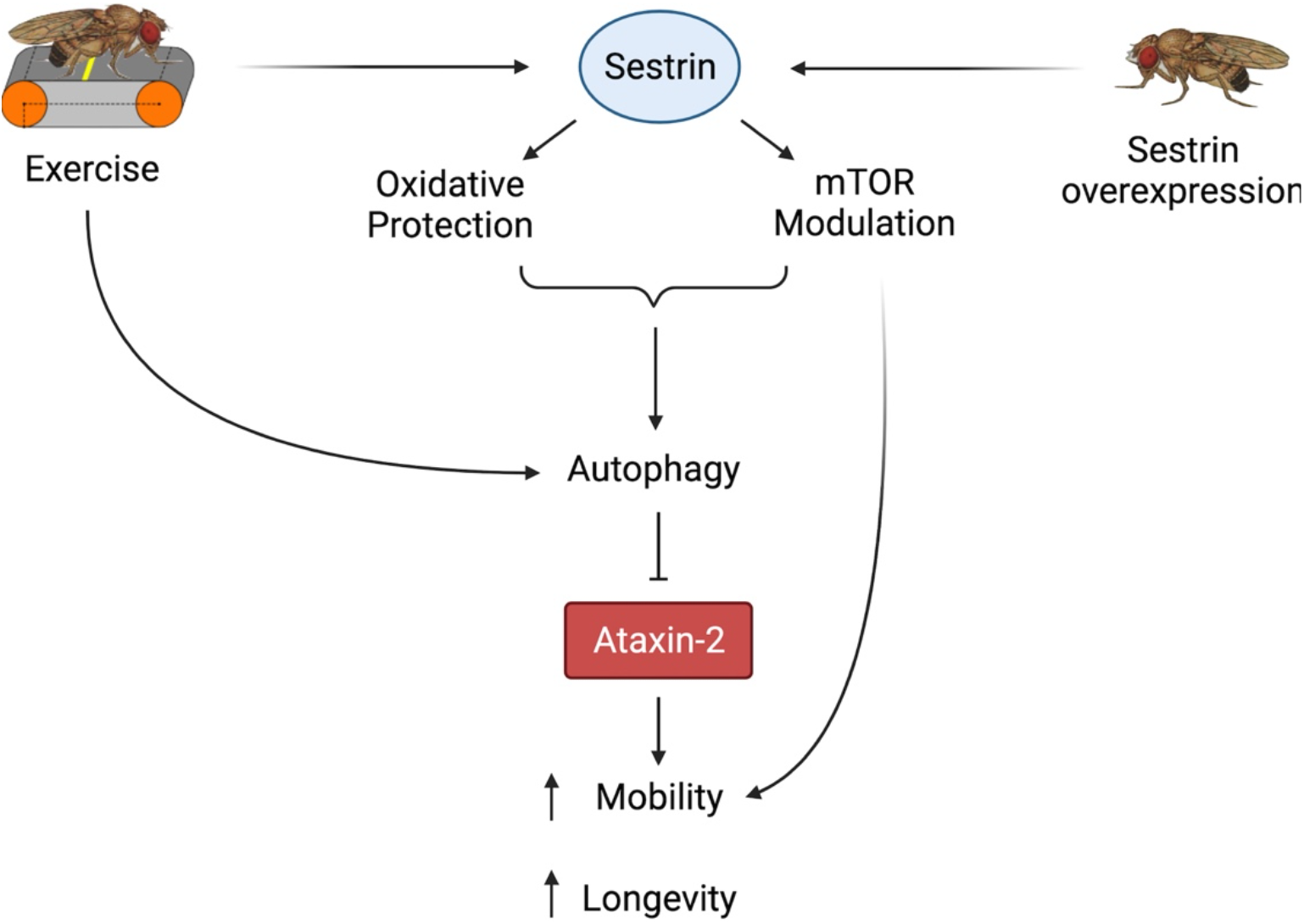
Proposed model of the effects of exercise and dSesn overexpression on SCA2 flies. Sestrin’s known functions and activation by exercise have been established previously (39,62). Wild type Sesn overexpression or exercise can activate autophagy and reduce disease protein levels, improving function.

In summary, both exercise and dSesn expression ameliorate mobility and survival defects in SCA2 model flies and these improvements correlate with reduced disease protein levels. Furthermore, these neuroprotective effects require known functions of dSesn related to mTOR and oxidoreductase activities.

## Discussion

Endurance exercise enhances quality of life and provides remedy for a number of conditions and diseases, but, until now, has not been systematically examined across several polyQ disorders. Lack of systematic studies in this respect also means that exercise-dependent molecular mechanisms that can be utilized to fight polyQ diseases remain largely unknown.

Here, we explored in a highly controlled and rigorously quantified way the possibility that daily training is of benefit against some polyQ diseases, but not necessarily others. In addition, we identified key mediators of exercise’s protective effects that can be harvested towards therapeutic options for individuals unable to exercise, or to enhance benefits in those who can.

Endurance exercise significantly rescued progressive motility defects in SCA2 flies, allowing their climbing speed and endurance to remain in wild-type range. Exercise also partially rescued SCA6 flies, with no beneficial effect on SCA3 flies. Thus, exercise can have potent rescue effects for particular polyQ diseases. To obtain early insight into how exercise could lead to such dramatic improvement in SCA2 flies (albeit not for SCA3) we examined protein levels of ataxin-2 and ataxin-3, their respective disease proteins. Exercise led to clear reduction in protein levels of ataxin-2, but did not noticeably affect ataxin-3. In the context of endurance exercise, improved motility in SCA2 flies may be due to reduced disease protein levels.

In addition, we examined the effect of the powerful exercise-mimetic protein Sesn on lifespan, motility and disease protein levels in SCA2 model flies. Sesns are induced by oxidative and genotoxic stress in flies and mammals and promote autophagy (58, 64). Sesns protect against oxidative stress-induced fly neuronal death (65) and reduce accumulation of aggregates in mammalian cells (66). In previous work with wild-type flies, we found that expression of fly *Sesn* is sufficient to preserve speed and endurance and has no additive effect with exercise, consistent with a role as a key effector of exercise (39). Here, we found that Sesn was protective in SCA2 model flies, but did not confer benefit to either SCA3 or SCA6. Furthermore, while Sesn’s oxidoreductase function was not necessary to rescue mobility defects in SCA2, both oxidative resistance and mTOR modulating activities were required for Sesn to improve early death. In fact, overexpression of Sesn with mutations eliminating either its oxidoreductase activity or mTOR modulating functions exacerbated premature death in SCA2 flies.

We have previously shown that the oxidoreductase activity of Sesn is dispensable for motility improvements in wild-type flies (39), similar to our observations with SCA2 flies in this study. Sesn overexpression is also not sufficient to extend lifespan in a wild-type background, shown here and elsewhere (39, 58). Neurons are particularly susceptible to oxidative stress and reactive oxygen species can induce cell death during neurodegeneration (67). Perhaps, in the context of neurodegenerative disease, Sesn protects mobility until oxidative burden in neurons becomes insurmountable.

Wild-type Sesn overexpression in SCA2 model flies correlates with significantly reduced disease protein and increased autophagic flux. Overexpression of mutated versions of Sesn led to reduced disease protein levels without reaching statistical significance. This is not entirely surprising: mutated Sesns still had some positive impact on SCA2 fly mobility, but did not have the same, powerful protective effect of wild-type Sesn. Additionally, mutant Sesns, as mentioned above, failed to improve lethality in SCA2 model flies. Collectively, these findings suggest separate mechanisms at play: some that may depend on reduced toxic protein levels (as observed before in a model of HD and exercise (68, 69)), and others that may protect independently of changes in degenerative protein levels. Future investigations are required to parse out such mechanisms and whether they function in a cell autonomous or non-cell autonomous manner.

To conclude, we propose endurance exercise as a promising therapeutic intervention for polyQ neurodegeneration, particularly in SCA2. We also identify Sesn as a genetic mediator of exercise-induced improvements in SCA2 phenotypes and demonstrate that Sesn can substitute for exercise benefits, even in flies that do not exercise. Identification of molecular targets capable of inducing exercise-like enhancements in the presence of neurodegeneration is particularly important, as many patients will eventually lose the ability to exercise. This work has begun to uncover exercise-induced mechanisms that can be utilized for neuroprotection against specific polyQ diseases and can be leveraged toward therapeutics in the future.

## Methods

### Antibodies

Primary antibodies were obtained from the following sources: anti-ataxin-2 (mouse monoclonal, 1:500, BD biosciences) anti-ataxin-3 (mouse monoclonal 1H9, MAB5360, 1:500-1000; Millipore), anti-MJD (rabbit polyclonal, 1:15,000)(70)), anti-dSesn (rabbit polyclonal, 1:500, (58)), anti-GABARAP (rabbit polyclonal, Abcam, 1:1000) Peroxidase conjugated secondary antibodies (goat anti-mouse, goat anti-rabbit, 1:5000; Jackson Immunoresearch).

### Fly Stocks and Maintenance

Gifted stocks used in this study were sqh-Gal4 (Daniel Kiehart, Duke University), and w^1118^ (Russ Finley, Wayne State University). UAS-Q80, UAS-ATXN3^Q77^, UAS-α1ACT^Q70^ and elav-Gal4 (GS) were previously described in (44, 54, 60) UAS-ATXN2^Q117^ (BDSC#68394, BDSC#68395), on chromosomes II and III, respectively, were obtained from the Bloomington *Drosophila* Stock Center (Bloomington, IN). Both SCA2 lines were used for pilot work. A single line containing UAS-ATXN2^Q117^ on both chromosomes II and III was then generated using standard crosses (w-;UAS-ATXN2^Q117^/CyO;UAS-ATXN2^Q117^).

Wild-type (dSesn^WT^) and C86S(dSesn^CS^), D424A (dSesn^DA^) or D423A/D424A (dSesn^DDAA^) mutations are described in (39, 62). C86, D423 and D424 in dSesn correspond to C125, D406 and D407 in mammalian SESN2 (62). These four lines were separately recombined with sqh-Gal4 to create four independent lines with genotypes: +;sqh/CyO;UAS-dSesn^XX^/TM3-Sb. Simultaneous overexpression of both UAS-ATXN2^Q117^ and UAS-dSesn^XX^ was achieved via a single cross between w-; UAS-ATXN2^Q117^/CyO;UAS-ATXN2^Q117^ virgin females and +;sqh/CyO;UAS-dSesn^XX^/TM3-Sb males to generate +;UAS-ATXN2^Q117^;UAS-dSesn^XX^/ATXN2^Q117^. All insertions were validated by Western blotting (42, 44, 46, 54, 58, 60, 71, 72).

Prior to all experiments, fly cultures were maintained at a constant density for at least two generations. 20-25 virgin females (depending on genotype) and 5 males were mated in 300 mL bottles with 50 mL standard 10% sucrose 10% yeast spiked with 500 μL Penicillin-Streptomycin (10,000 u/mL, 10mg/mL in 0.9% sterile NaCl (Sigma Aldrich). Adult progeny were synchronized by collecting within 6 hours of eclosion over a 24 hour time period. Groups of 20 age and sex-matched flies were immediately transferred into narrow polypropylene vials containing 5mL of standard 10% sucrose 10% yeast (no antibiotic) or RU486 food as appropriate. Food vials were changed every day during exercise and every second day thereafter and scored for mobility and lifespan until no flies remain.

Flies were housed in a 25°C incubator on a 12:12 light:dark cycle at 50% relative humidity. Control flies for all non-gene-switch Gal4 UAS experiments consisted of both the UAS and Gal4 lines into *w^1118^*. For gene-switch experiments, RU-flies of the same genotype served as the negative control. RU+ group received 100 μM RU486/mifepristone (Cayman Chemical, Ann Arbor, MI), which activates the gene switch (GS) driver, while RU-group received the same volume of vehicle solution (70% ethanol).

### Exercise Training

Triplicate cohorts of at least 800 flies were collected under light CO_2_ anesthesia within 6 hours of eclosion and separated into vials of 20. Flies were then further separated into 2 large cohorts of at least 400 flies, which served as exercised and unexercised groups. Exercised groups received three weeks of ramped exercise as described (29). The unexercised groups were placed on the exercise training device at the same time as the exercised groups but were prevented from running by the placement of a foam stopper low in the vial. Both cohorts were housed in the same incubator with normal foam stopper placement at all times other than during an exercise bout.

### Climbing Speed

Negative geotaxis was assessed in Rapid Negative Geotaxis (RING) assays in groups of at least 100 flies as described (73). Briefly, vials of 20 flies were briskly tapped down, then measured for climbing distance after 2s of inducing the negative geotaxis instinct. For each group of vials, an average of 5 consecutive trials was calculated and batch processed using ImageJ (Bethesda, MD). Flies were longitudinally tested 2-3 times per week for 5-7 weeks. Between assessments, flies were returned to food vials and housed normally as described above. Negative geotaxis results were analyzed using two-way ANOVA analysis (age-effect, genotype effect) with post hoc Tukey multiple comparison tests in GraphPad Prism (San Diego, CA, USA). All negative geotaxis experiments were performed in triplicate, with one complete trial shown in each graph.

### Endurance

At least eight vials of 20 flies from each cohort were subjected to the endurance analysis on day 25 of adulthood, immediately after exercise was complete and after 2 days of recovery. For each session, the flies were placed on the Power Tower exercise machine (28) and the climbing instinct was induced until flies no longer responded to the negative geotaxis stimulus. Monitored at 15 min intervals, a vial of flies was visually determined to be fatigued when 10% or fewer flies could climb higher than 1 cm after three consecutive drops. Each vial was plotted as a single datum. Endurance experiments were performed in triplicate and at the same time as age-matched background controls and were scored blindly when possible. The time from the start of the assay to the time of fatigue was recorded for each vial, and the data analyzed using log-rank analysis in GraphPad Prism (San Diego, CA, USA). Each graph represents an independent, representative repetition.

### Longevity

For exercise experiments, appropriate food vials (RU486, vehicle or standard 10% sucrose 10% yeast) were changed and deaths were recorded five times per week during exercise training and three times per week after completion of the training program. All other longevity experiments were scored three times per week. Dead flies were removed and counted until no flies remained. Differences in survival were plotted as probability of death and assessed following censoring using log-rank analysis in GraphPad Prism. Longevity experiments were performed in triplicate and in parallel with background controls, with each individual graph depicting a representative biological repetition.

### Western Blots

Three to five whole flies per biological replicate, depending on experiment, were homogenized in boiling lysis buffer (50 mM Tris pH 6.8, 2% SDS, 10% glycerol, 100 mM dithiothreitol), sonicated, boiled for 10 min, and centrifuged at 13,300xg at room temperature for 10 min. Western blots were developed ChemiDoc (Bio-Rad) and quantified with ImageLab (Bio-Rad). For direct blue staining, PVDF membranes were submerged for 10 min in 0.008% Direct Blue 71 (Sigma-Aldrich) in 40% ethanol and 10% acetic acid, rinsed in 40% ethanol/10% acetic acid, air dried, and imaged. Western blots were performed using at least 3 biological replicates, and statistical analysis was performed in GraphPad Prism.

### Statistical Analysis

Survival and endurance analyses were analyzed by log-rank. Non-natural deaths were censored and did not exceed 10% of flies for any group. Climbing speed over time was analyzed by Two-way ANOVA for age effect and genotype effect, with Tukey post-hoc comparison between significantly different groups. Western blots were analyzed using either student t-test or ANOVA depending on experiment. All statistics were performed using GraphPad Prism software version 9.2.0 for Macintosh.

## Acknowledgements

We thank Henry Paulson for the anti-MJD antibody, and Todi lab members Kozeta Libohova, Wei-Ling Tso, Gorica Ristic, Joanna Sutton, Jessica Blount, and Sean Johnson for generation and validation of the SCA lines used in this study. We acknowledge our collaborators Myungjin Kim and Jun Hee Lee for the original dSesn *Drosophila* stocks and reagents.

## Supplementary Figure legends

**Supplementary Video SV1: SCA2 model flies increase motility with exercise.** Video of 4-week-old uninduced control and SCA2 model flies taken following 3 weeks of ramped endurance exercise. Order of vials depicted, left to right: Vial 1-uninduced, exercised control, Vials 2,3-SCA2 model, exercised, Vials 4,5-SCA2 model, unexercised.

**Supplementary Figure S1:**
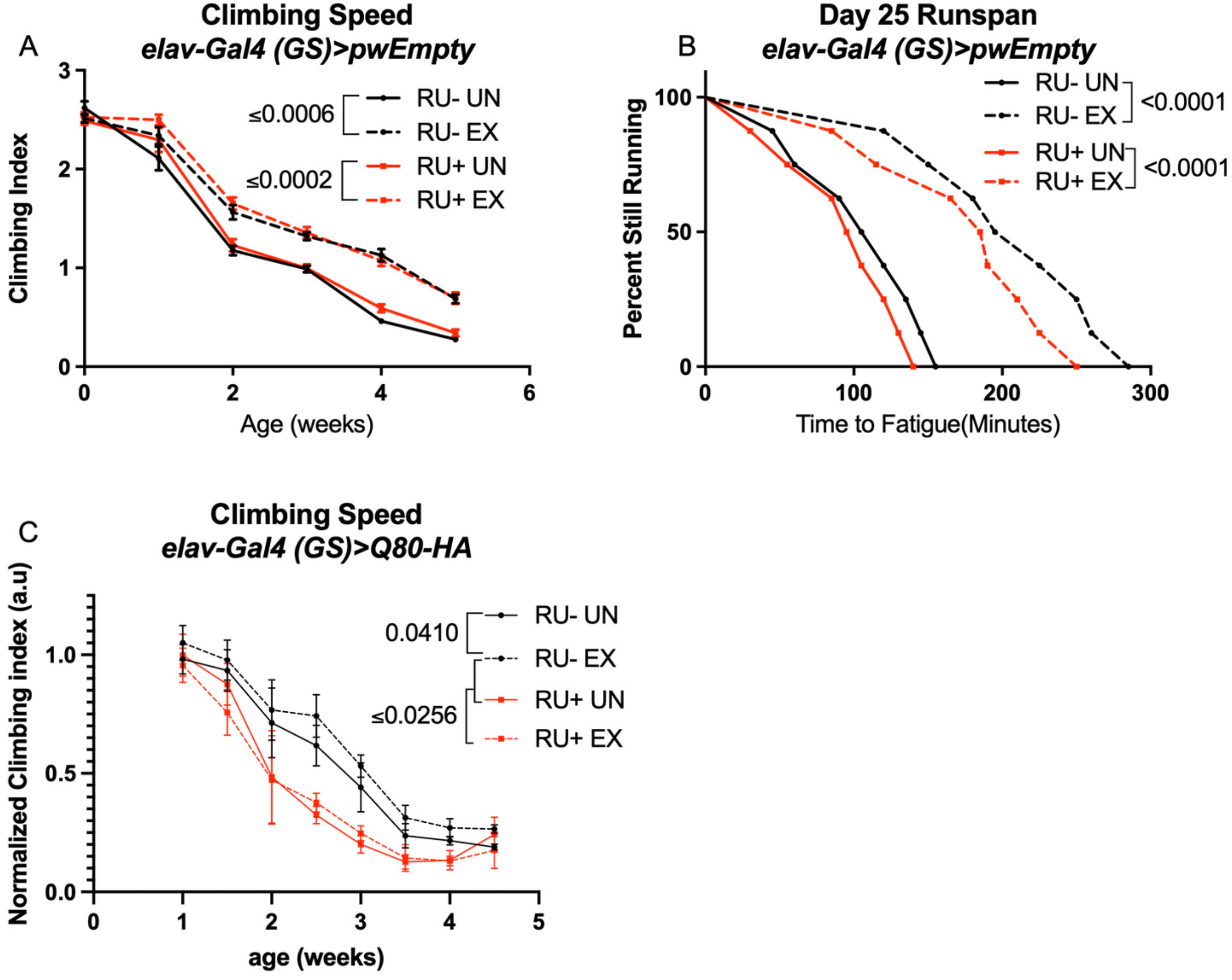
Endurance exercise improves climbing speed in uninduced background control flies. **(A, B)** *elav-Gal4 (GS)>pwEmpty* vector flies improve **(A)** climbing speed and **(B)** endurance independent of RU486-feeding. **(C)** Flies expressing an isolated polyQ-80 repeat in adult neurons have lower climbing speed than age-matched, uninduced control flies and do not increase speed with exercise. Mobility and endurance experiments performed in triplicate. n≥100 (climbing speed), n≥8 vials of 20 flies (endurance). Error bars indicate +/-SD.

**Supplementary Figure S2:**
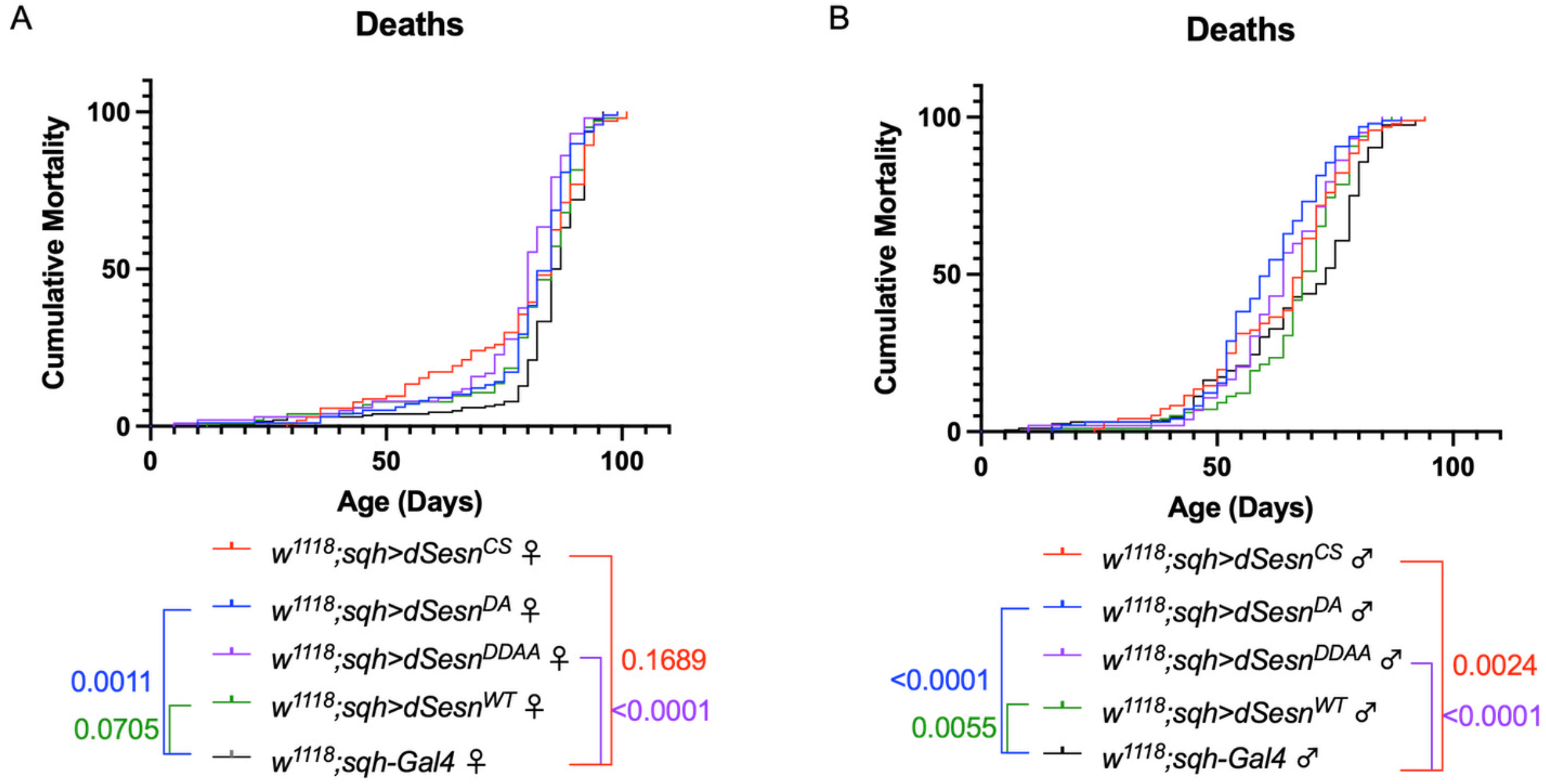
Ubiquitous dSesn expression in wild-type flies does not account for survival differences observed in flies expressing polyQ-expanded ATXN2. Survival curves in female **(A)** and male **(B)** flies expressing either wild-type dSesn (green), dSesn^CS^ (red), dSesn^DA^ (blue), dSesn^DDAA^ (purple) or background control (black). Flies ubiquitously expressing dSesn mutations have similar or lower survival than flies ubiquitously expressing wild type dSesn. n≥97.

## Source data-all figures

Tab delimited excel file containing raw data for all figures and supplementary figures. Uncropped, unedited Western blot data are labeled and included as separate jpegs.

